# A Rapid Electromechanical Model To Predict Reverse Remodeling Following Cardiac Resynchronization Therapy

**DOI:** 10.1101/2020.11.16.385161

**Authors:** Pim J.A. Oomen, Thien-Khoi N. Phung, Kenneth C. Bilchick, Jeffrey W. Holmes

**Affiliations:** Department of Biomedical Engineering, University of Virginia, Box 800759, Health System, Charlottesville, VA 22903, USA; Department of Environmental Health, Harvard T.H. Chan School of Public Health, 665 Huntington Ave, Boston, MA 02115, USA; Department of Medicine, University of Virginia, Box 800158, Health System, Charlottesville, VA 22903, USA; Department of Biomedical Engineering, University of Virginia, Box 800759, Health System, Charlottesville, VA 22903, USA School of Engineering, University of Alabama at Birmingham, 1075 13th St S, Birmingham, AL 35233, USA

**Keywords:** Growth, Hypertrophy, Heart failure, Cardiac Resynchronization Therapy, Dyssynchrony, Patient-specific modeling

## Abstract

Cardiac resynchronization therapy (CRT) is an effective therapy for patients who suffer from heart failure and ventricular dyssynchrony such as left bundle branch block (LBBB). When it works, it reverses adverse left ventricular (LV) remodeling and the progression of heart failure. However, CRT response rate is currently as low as 50-65%. In theory, CRT outcome could be improved by allowing clinicians to tailor the therapy through patient-specific lead locations, timing, and/or pacing protocol. However, this also presents a dilemma: there are far too many possible strategies to test during the implantation surgery. Computational models could address this dilemma by predicting remodeling outcomes for each patient before the surgery takes place. Therefore, the goal of this study was to develop a rapid computational model to predict reverse LV remodeling following CRT. We adapted our recently developed computational model of LV remodeling to simulate the mechanics of ventricular dyssynchrony and added a rapid electrical model to predict electrical activation timing. The model was calibrated to quantitatively match changes in hemodynamics and global and local LV wall mass from a canine study of LBBB and CRT. The calibrated model was used to investigate the influence of LV lead location and ischemia on CRT remodeling outcome. Our model results suggest that remodeling outcome varies with both lead location and ischemia location, and does not always correlate with short-term improvement in QRS duration. The results and time frame required to customize and run this model suggest promise for this approach in a clinical setting.

## 1 Introduction

Heart failure is associated with 300,000 deaths annually in America alone, while over a million Americans experience a myocardial infarction each year (Mozaffarian et al. 2015). Many patients with heart failure and/or myocardial infarction also develop left ventricular dyssynchrony, which exacerbates cardiac dysfunction, worsens symptoms and decreases survival (Baldasseroni et al. 2002). Left bundle branch block (LBBB), in which there is abnormal conduction on the left ventricular septum, (Bilchick et al. 2007; Shamim et al. 1999) leads to mechanical dyssynchrony characterized by delayed contraction and increased stretching of the LV free wall relative to the interventricular septum. (Prinzen et al. 1995; Vernooy et al. 2005; Aalen et al. 2019; Fixsen et al. 2019). As a consequence, LV pump function is impaired, and chronic LV dilation and asymmetric wall thickening occur (Baldasseroni et al. 2002; Vernooy et al. 2005; Prinzen et al. 1995). Ventricular dilation exacerbates mechanical dyssynchrony, which, in turn, promotes further impairment of LV function and further LV dilation in a vicious cycle (Kerckhoffs et al. 2012b).

Cardiac resynchronization therapy (CRT) has emerged as an effective therapy for many patients who suffer from heart failure and left ventricular dyssynchrony. CRT is implemented through a pacemaker designed to restore coordinated contraction of the heart by pacing from multiple locations at appropriate times (Bilchick et al. 2007). When CRT is successful, it improves survival by stopping and even reversing LV remodeling (Bristow et al. 2004; Cleland et al. 2005; St. John Sutton et al. 2003; Vernooy et al. 2007); however, 35-50% of patients do not derive sufficient benefit from CRT to be classified as CRT responders(Chung et al. 2008; Tracy et al. 2012). The likely reason for this observation is that the long-term remodeling response to stimulation from any one left ventricular pacing location with CRT varies significantly from patient to patient based on mechanical activation time, scar size, and scar location (Bilchick et al. 2014; Rademakers et al. 2010). As a result, a personalized rather than a uniform approach to CRT is expected to be optimal. Specifically, CRT should account for patient-to-patient variability by allowing clinicians to tailor the therapy through the use of patient-specific lead locations, timing intervals for pacing, and choice of quadripolar lead vectors. Unfortunately, in practice there are far too many possible strategies to test during the implantation surgery. As a result, there is a critical need for computational models that can predict CRT remodeling outcomes before the surgery takes place.

Current computational models that aim to guide CRT focus primarily on acute outcomes such as mechanical dyssynchrony and left ventricular pump function (Niederer et al. 2011; Walmsley et al. 2015; Huntjens et al. 2014; Constantino et al. 2012). While it seems plausible that better initial mechanical synchrony or function should lead to better outcomes, these models do not necessarily predict long-term LV remodeling that ultimately determine CRT success. Recently, it has become possibe to make accurate predictions for these long-term outcomes, as multiple computational models have utilized changes in strain to predict observed trends in cardiac remodeling in response to pressure and volume overload (Arts et al. 2005; Genet et al. 2016; Göktepe et al. 2010; Kerckhoffs et al. 2012a; Kroon et al. 2009) in the setting of LBBB (Kerckhoffs et al. 2012b; Arumugam et al. 2019). These models are typically implemented in a finite-element framework, and many of them require hours or days to customize for an individual patient in order to simulate a predicted time course of remodeling for a single disease or intervention. Moreover, most models have only addressed adverse ventricular remodeling such as resulting from heart failure with LBBB rather than favorable reverse remodeling following an intervention such as CRT.

Thus, the goal of this study was to develop a rapid computational model to predict reverse LV remodeling following CRT. We recently developed a rapid computational model that predicts LV remodeling following volume overload, pressure overload, and myocardial infarction based on changes in myocardial strain (Witzenburg and Holmes 2018). Here, we adapted this model to simulate the mechanics of ventricular dyssynchrony, added a rapid electrical model to predict electrical activation timing, and implemented an evolving homeostatic setpoint in the growth law, which we recently showed can be useful in capturing reverse remodeling (Yoshida et al. 2019). The model was calibrated to published experimental data and applied to predict the influence of CRT lead placement on LV remodeling.

## 2 Methods

The model code employed here is available for download from GitHub (https://github.com/PimOomen/CardioGrowth, https://github.com/tkphung/CardiacElectricalActivation).

### 2.1 Electrical model for ventricular dyssynchrony

The electrical model consisted of a biventricular finite-element mesh parameterized by material properties describing electrical conduction velocity in the elements. The geometry of the finite-element mesh was generated from a canine-specific cardiac MRI using the open-source method described by Phung et al. (2019). The material properties in the elements include the myocardial fiber orientation and conduction velocities. To describe the myocardial fiber architecture, we used a rule-based method to assign fibers with orientations varying linearly from 60 on the endocardium to −60 at the epicardium in the plane of the wall (Streeter and Hanna 1973). For each element, the conduction velocities were assigned along the fiber orientation, transmurally through the wall, and in the cross-fiber orientation in the plane of the wall. The heart was divided into a fast, endocardial layer in the right and left ventricles and the remaining bulk myocardium. Conduction speed was assumed to be transversely anisotropic with faster conduction along the fiber direction and 40% of that velocity in the cross-fiber directions. We specified endocardial fiber velocity to be 600% of the bulk myocardium fiber velocity to model the Purkinje network as a fast conduction layer (Aguado-Sierra et al. 2011; Hyde et al. 2015; Kayvanpour et al. 2015; Lee et al. 2019; Okada et al. 2017; Hunter and Smaill 1988). Absolute conduction velocities were scaled by changing the bulk myocardium fiber velocity while maintaining the same scaling for the cross-fiber and endocardial fiber velocities (Lee et al. 2019). To simulate ischemia, we mapped elements to an ischemic region and removed them from the electrical simulation to represent complete electrical block.

We simulated electrical wavefront propagation using a cellular automata framework developed by Hunter and Smaill (1988). The model is initialized by specifying an element from which the electrical wavefront starts. The framework relies on each element having a discrete state, which we specify to be resting or depolarized. The activation starts from a depolarized initiation element and spreads to resting neighboring elements at a time interval dependent on the conduction velocities and distances between the elements centroids. Because we have discretized the geometry into finite elements with defined neighbors (touching elements), we represent the model as a weighted, undirected graph. The nodes of the graph are the centroids of all elements. The edges are the connections between each element and its neighbors. The weights of the edges are calculated as the time required for the wave front to pass from the centroid of an element to its neighbor. After specifying an initiation element, we can solve the propagation efficiently in MATLAB (Natick, MA) using a shortestpath tree algorithm emanating from the initiation site. The algorithm connects the initiation node to every other node by traversing the shortest combination of edges. The result of the cellular automata simulation is the electrical activation time for each element in the biventricular mesh. To couple the output from the electrical model with the compartmental model, we divided the left ventricular finite-element mesh and averaged the electrical activation times in 16-segment AHA regions. The right ventricular elements were averaged in 5 segments (3 basal and 2 apical).

To compare the electrical model to experimental data, 12-lead pseudo-ECG signals were calculated from the simulated electrical activation times using a formulation based on Weinberg et al. (2008); Roberts et al. (1979); Miller and Geselowitz (1978). Briefly, we set the transmembrane potential for each element to be a Heaviside step function which steps from −90 mV to 0 mV at the electrical activation time. We calculated the gradient of the transmembrane potential across the biventricular geometry and projected the contribution of each element to a particular ECG lead. Because the transmembrane potential step function only simulates depolarization, the pseudo-ECG simulates the lead voltage for an equivalent depolarization phase for comparison to the ECG data.

### 2.2 Compartmental model of the circulation

We used our previously published lumped-parameter model of the circulation to account for hemodynamic loading. Systemic and pulmonary arteries and vessels were modeled as capacitators in parallel with resistors, and heart valves as pressure-sensitive diodes, as described previously (Fig. 1) (Witzenburg and Holmes 2018; Santamore and Burkhoff 1991). The blood volume contained in each capacitive compartment was functionally divided into two parts: unstressed and stressed blood volume. The unstressed blood volume was defined as the maximum blood volume that can fit within a compartment without causing the pressure to rise from 0 mmHg, while any blood volume in excess of that was defined as the stressed blood volume. Here we assumed the unstressed blood volumes of compartments remained constant, while stressed blood volume could change between compartments. Systemic vascular resistance (SVR) and SBV were adjusted to match reported baseline and acute hemodynamics (Section 2.5). All other resistances and capacitances were adopted from Santamore and Burkhoff (1991). The model was implemented in MATLAB as a series of differential equations for changes in volume of each compartment (LV, systemic arteries, systemic veins, RV, pulmonary arteries, and pulmonary veins).

**Fig. 1.**
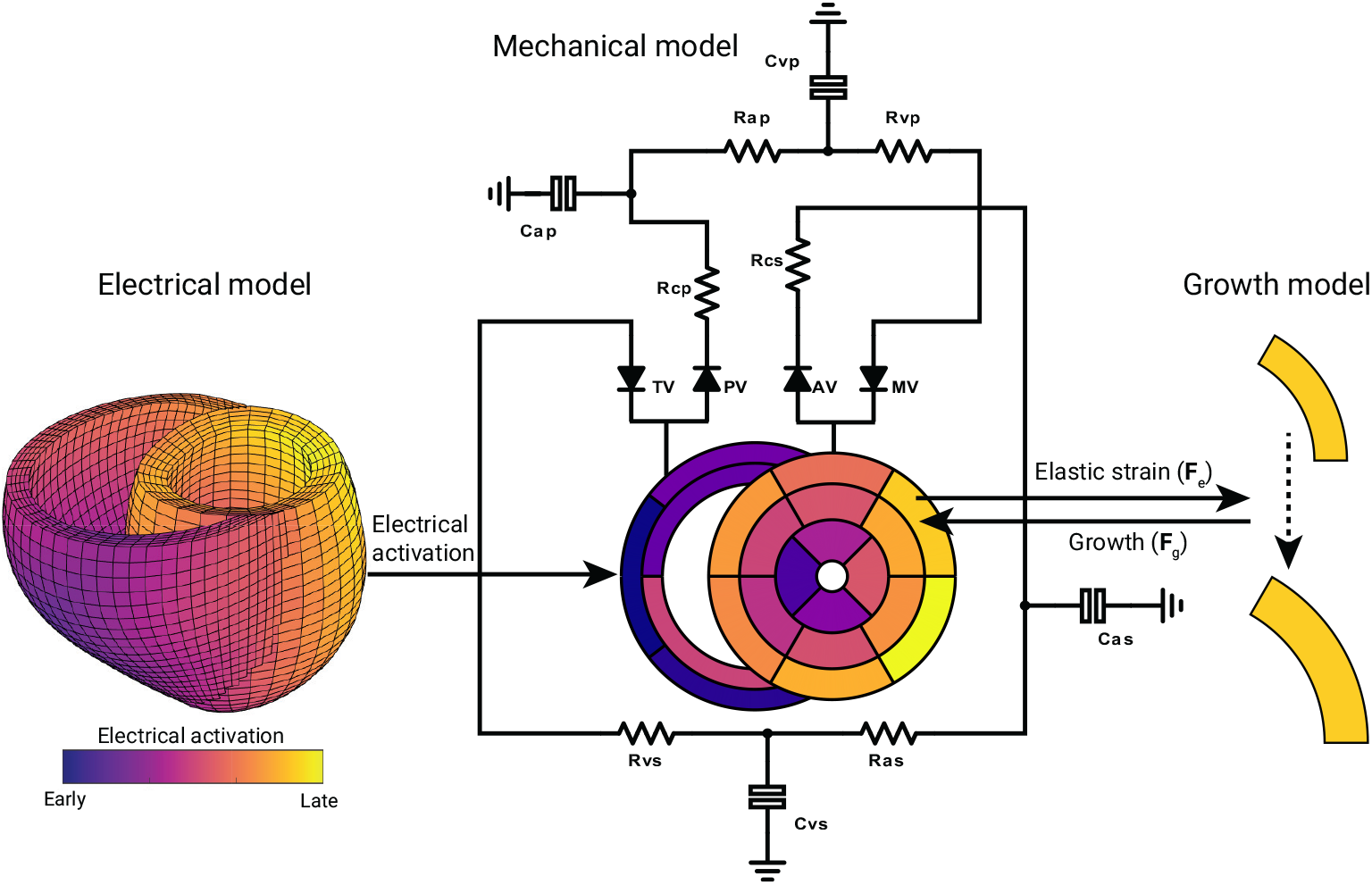
Schematic representation of the electromechanical model. A graph-based finite element model was used to calculate the electrical activation timing through the left and right ventricle. Electrical activation timing was averaged in 16-segment AHA regions (and 5 RV segments) for coupling to the simplified model of the heart (Lumens et al. 2009; Walmsley et al. 2015) and circulation (Witzenburg and Holmes 2018). Systemic and pulmonary arteries and vessels were modeled as capacitators in parallel with resistors, and heart valves as pressure-sensitive diodes. Remodeling of each segment was determined using an isotropic strain-driven growth model with evolving setpoint.

### 2.3 Simplified mechanical model of the heart

In our previous work, we modeled each ventricular compartment as a thin-walled sphere, with no direct interaction between the LV and RVWitzenburg and Holmes (2019). To account for intraventricular differences in electrical activation and wall strain, as well as LV-RV interaction, we here adopted the TriSeg approach from (Lumens et al. 2009). The ventricular geometry was simulated as three thick-walled spherical segments: left free wall (LFW), right free wall (RFW), and septal wall (SW). The spherical walls encapsulated the two ventricular cavities, and coincided at a junction circle. The ventricular geometry was assumed to be axisymmetric around an axis perpendicular to this junction circle. Using the MultiPatch approach from (Walmsley et al. 2015), the LFW and SW were divided into segments according to the 16-segment AHA model (11 segments for the LFW, 5 segments for the SW), while the RFW was divided into 5 segments (Fig. 1). All segments within one wall share the same curvature at any time point during the cardiac cycle, but can have different stiffnesses due to local differences in electrical activation timing (determined by the electrical model, Section 2.1). Consequently, stresses and strains can differ between segments in the same wall.

Mechanics were calculated for each segment at the mid-wall plane *A*_m,w_ (w = LFW, RFW, SW), which was defined as the spherical surface that divides wall volume (*V*_w_) in in-ner and outer parts of equal volume. We assumed the myocardium to be incompressible, with myofibers distributed isotropically along the midwall plane. Passive midwall Cauchy fiber stress *σ*_ff,pas_ was modeled using an exponential consti-tutive model:

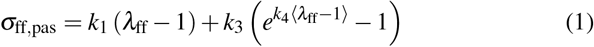

Where *λ*_ff_ is segmental elastic fiber stretch and *k*_1_, *k*_3_, and *k*_4_ material parameters. Midwall active stress (*σ*_ff,act_) was cal-culated using a Hill-type sarcomere active contraction model (Lumens et al. 2009; Walmsley et al. 2015), in brief:

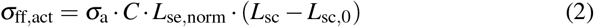

where *σ*_a_ is a constant that describes the maximum stress the muscle can generate; C is a state variable of segmental muscle contractility as a function of time and fiber stretch; *L*_se,norm_ is the length of the normalized series elastic element in the Hill-type model; and *L*_sc_ and *L*_sc,0_ are the sarcomere contractile element length at current and unloaded state, respectively. Onset of contraction (*C* > 0) for each segment was obtained from the electrical model (Section 2.1). From the midwall Cauchy fiber stress in each segment, midwall segment tension *T*_s_ was calculated by principle of conservation of energy (Lumens et al. 2009):

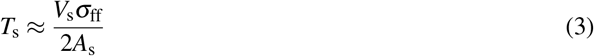

The tension of each wall was then estimated using the MultiPatch method, which utilizes a linearization of the wall tension around a working point corresponding to the wall at zero wall tension (Walmsley et al. 2015). An iterative Newton scheme was employed to achieve balance of tension of the three walls at the junction point. This was achieved by adjusting the radius of the junction circle and position of the SW relative to the LFW and RFW until balance of tension converged under 1e-6 N.

### 2.4 Strain-based growth model with evolving setpoint

Myocardial growth was simulated using a kinematic growth approach, where the total deformation gradient tensor ***F*** is multiplicatively decomposed into an elastic contribution ***F***_e_ and a growth contribution ***F***_g_:

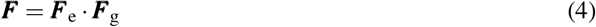

Changes in myocardial strain were used to determine the amount of cardiomyocyte growth. Previously in our model, cardiomyocyte lengthening was determined by changes in maximum elastic fiber Green-Lagrange strain *E*_ff_, while thickening was determined by the change in maximum elastic radial strain *E*_rr_ (Witzenburg and Holmes 2018). To match the growth pattern typically observed during LBBB, we here imposed isotropic growth that was driven by changes in maximum segmental *E*_ff_:

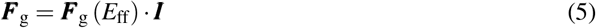

with ***I*** the second-order identity tensor. At each growth time step *i*, the so-called growth stimulus was calculated for each LV segment:

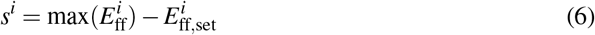

where 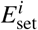 is a homeostatic setpoint, which was initially defined as the maximum fiber Green-Lagrange strain at base-line, 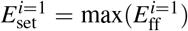, such that in a baseline simulation *s*^*i*=1^ = 0 and no growth would occur. After baseline, the homeostatic setpoint gradually changed during growth, an adaptation shown critical to predicting reverse growth following relief of pressure overload using mechanics-based growth laws by Yoshida et al. (2019). This was implemented using a weighted moving average of the previous elastic strains to specify the current setpoint. Strains were weighted using a triangular moving window spanning 45 days, where 23 days prior to the current growth time step *i* were weighed most heavily, and strains from more recent and distant time steps exerting less influence:

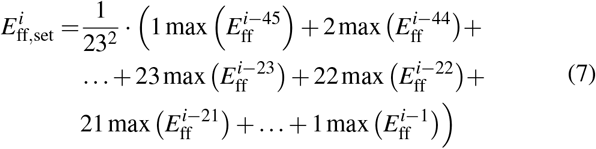

The stimulus function was used to define incremental growth via a sigmoidal relationship, with a quiescent zone around *s*^*i*^ = 0,

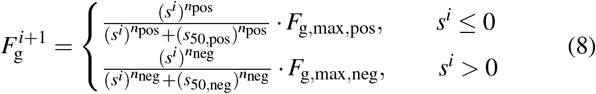

where *s*_50,pos_, *s*_50,pos_, *n*_pos_, and *n*_neg_ determine the shape of the sigmoid curve, and *F*_g,max,pos_ and *F*_g,max,neg_ define the maximum positive and negative growth rates. Growth parameter values can be found in the Supplement.

After each growth time step, the segmental wall volume and surface area were updated. Subsequently, the circulation model was solved using the 4th order Runge-Kutta method until a steady-state solution was established, i.e. the compartmental volumes at the beginning and end of the cardiac cycle were within 0.1 mL of each other. Model pseudocode is included in the Supplement

### 2.5 Model fitting

To calibrate our baseline electrical propagation model, we used recorded ECG data from six canines during surgery to induce LBBB. 12-lead ECGs were recorded before and immediately after radiofrequency ablation of the left bundle branch. Myocardial infarction was created in two out of the six canines one week prior to ablation by ligating of the left anterior descending coronary artery. One week after the induction of LBBB, we recorded Cine MRI to provide the biventricular geometry for the electrical model. The 9 ECG electrode locations were approximated on the torso geometry obtained from Cine MRI. We calibrated the model using a two-step approach similar to Giffard-Roisin et al. (2017). We optimized the intrinsic initiation element by simulating propagation from each endocardial element and comparing the subsequent pseudo-ECG to the recorded ECG. We normalized both signals ranges to focus on the direction and timing of the deflections rather than the magnitudes. We calculated the correlation coefficient between the predicted and recorded normalized ECGs to obtain a metric describing how well the shapes of the signals matched, and averaged the correlation coefficients of the 12 leads to obtain a single error metric for each simulation. Using the best fit initiation element, we scaled the myocardial fiber velocity to match the latest electrical activation time with the recorded QRS duration. To predict the effect of CRT, we introduced additional initiation elements in the model to represent CRT pacing leads. The pacing initiation elements were coordinated to depolarize simultaneously with the intrinsic initiation element.

To calibrate our growth model, we used experimental data published by Vernooy et al. (2005, 2007). In this study, LBBB was induced in canines via radiofrequency ablation, and biventricular CRT pacing was started after 8 weeks. Throughout the 16-week study, hemodynamic and growth data were collected. Since the growth model is driven by changes in strain, we first calibrated our pre and post-LBBB circulation and LV parameters using our previously published fitting method (Witzenburg and Holmes 2018). Model parameters at pre and acute post-LBBB were estimated by minimizing an objective function consisting of Z-scores of hemodynamic and kinematic measures. The experimental data we matched were: enddiastolic pressure (EDP), end-systolic pressure (ESP), end-diastolic volume (EDV), ejection fraction (EF, from Vernooy et al. (2005)), maximum rise in pressure (dp/dt_max_) and midwall segmental strain. EDP, ESP, dp/dtmax, and strain were matched for both pre- and post-LBBB. To compare strain data, the Pearson correlation coefficient between experimental and simulated midwall segmental strain traces was calculated. The objective function was minimized by adjusting the following model parameters using MATLABs fminsearch algorithm: circulation parameters (SBV and SVR, pre and post-LBBB), constitutive parameters (*k*_1_, *k*_3_, *σ*_a_), and geometry (unloaded midwall area). Additionally, LFW, RFW, and SW masses were derived from the same Cine MRI that was used to provide the biventricular geometry for the electrical model, and were scaled such that the total wall mass was equal to the reported wall mass at baseline in Vernooy et al. (2005).

After setting up the baseline and acute states of the model, we simulated the study protocol of Vernooy et al. (2007), i.e. 8 weeks of LBBB followed by 8 weeks of CRT with lateral midwall pacing. The growth parameters 8 were adjusted to match experimental changes of EDV, global wall mass, and lateral and septal wall mass. While adjusting the growth parameters, experimental EDP and ESP were matched at chronic (8 weeks) LBBB, acute post-CRT, and chronic (8 weeks) CRT by adjusting SVR and SBV.

### 2.6 Model predictions

Activation timing and ischemia at the LV lead are associated with CRT outcome (Bilchick et al. 2014; Rademakers et al. 2010). For non-ischemic patients, pacing at the site of latest electrical activation is believed to provide the best outcome, however we hypothesize that in ischemic LBBB patients, presence of scar can alter the relationship between optimal pacing site and pre-CRT activation. Therefore, we investigated the influence of lead location and scar pattern on post-CRT growth. Seven different cases were simulated: one non-ischemic LBBB case (using the model setup from the fitting study), and six ischemic LBBB cases. The ischemic cases consisted of ischemia patterns that can occur after basal or midwall occlusion of each of the main coronary arteries: left anterior descending artery (LAD), left circumflex artery (LCX), and right coronary artery (RCA). Ischemic regions were modeled as non-conductive (electrical model), and uncapable of contracting or growth (compartmental growth model). Growth was simulated for 8 weeks of LBBB and 8 weeks of CRT, without adjusting hemodynamic parameters since to the best of our knowledge no experimental measurements are available. For each case, influence of LV lead location on post-CRT growth was investigated by simulating the lead position in each one of the 16 segments, so in total 16 growth simulations were performed for each of the seven cases.

## 3 Results

### 3.1 Electrical activation model can simulate pre- and post-LBBB ECG data

We successfully fitted our electrical model to match pre- and post-LBBB 12 lead ECG data of 6 canines (4 non-ischemic, 2 ischemic). A comparison of all cases is included in Supplementary Figure 1; the case (C6) used as input for all the growth simulations in this study is shown in Fig. 2. The LBBB and baseline models differed in their initiation elements and myocardial fiber conduction velocities. The initiation element in the pre-LBBB model was in the basal RV free wall and moved apically towards the middle of the RV free wall for post-LBBB. Additionally, the myocardial fiber conduction velocity was slowed from 1.85 to 0.85 m/s to match the recorded QRS durations of 78 and 151 ms, respectively.

**Fig. 2.**
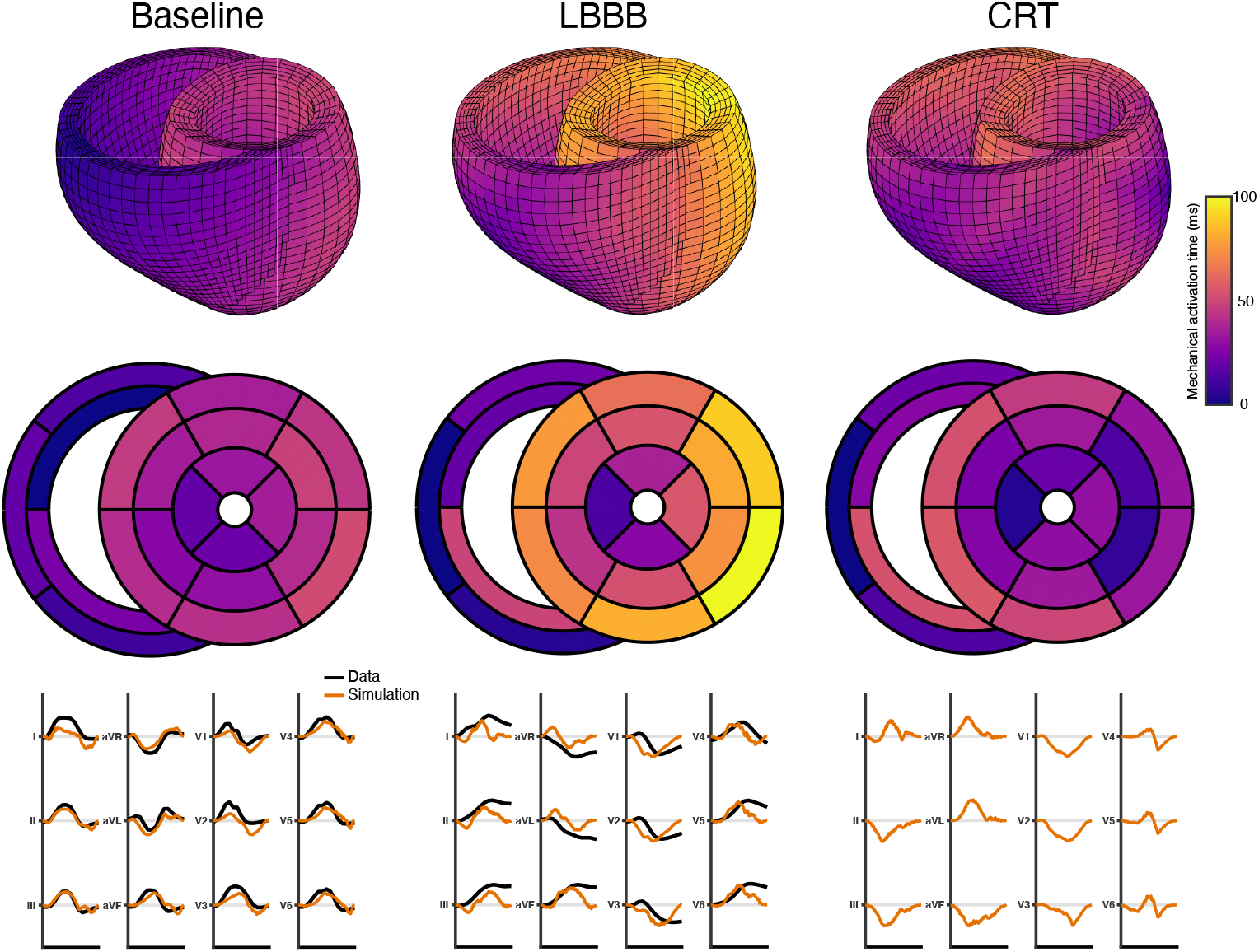
The baseline and left bundle branch block (LBBB) electrical simulations were fit to 12 lead ECG data obtained pre- and post-LBBB ablation surgery in a canine. The initiation site of electrical activation and the conduction velocity were optimized to match the QRS complex of the recorded 12 lead ECG data. CRT was simulated by adding an additional electrical initiation site (here in the LV lateral midwall). The electrical activation times were averaged into the 16-segment AHA regions for the LV and 5 regions for the RV.

### 3.2 Rapid computational model can simulate reverse remodeling following CRT using changes in strain

We successfully fitted our mechanical model of the heart and circulation to pre- and post-LBBB experimental data (Vernooy et al. 2007), and adjusted our growth parameters to match post-LBBB and post-CRT growth. 16 weeks of growth following LBBB and CRT were simulated in just under one minute on a laptop computer (8 GB RAM, 64-bit operating system, 2.9 GHz Dual-Core Intel Core i5).

Mechanical dyssynchrony increased acutely after LBBB (Fig. 3a,b). Maximum fiber strain, which drives growth in our model (Eq. 6), increased in most lateral wall segments. In contrast, no clear difference in maximum strain was observed in the septal wall segments. During baseline, maximum fiber strain was reached at the end of isovolumetric contraction, whereas after LBBB this occurred in the initial phase of ejection. After simulating CRT with lateral midwall pacing, mechanical synchrony was reestablished and maximum strain decreased in the lateral wall, however an increase in maximum strain occurred in the septal wall.

**Fig. 3.**
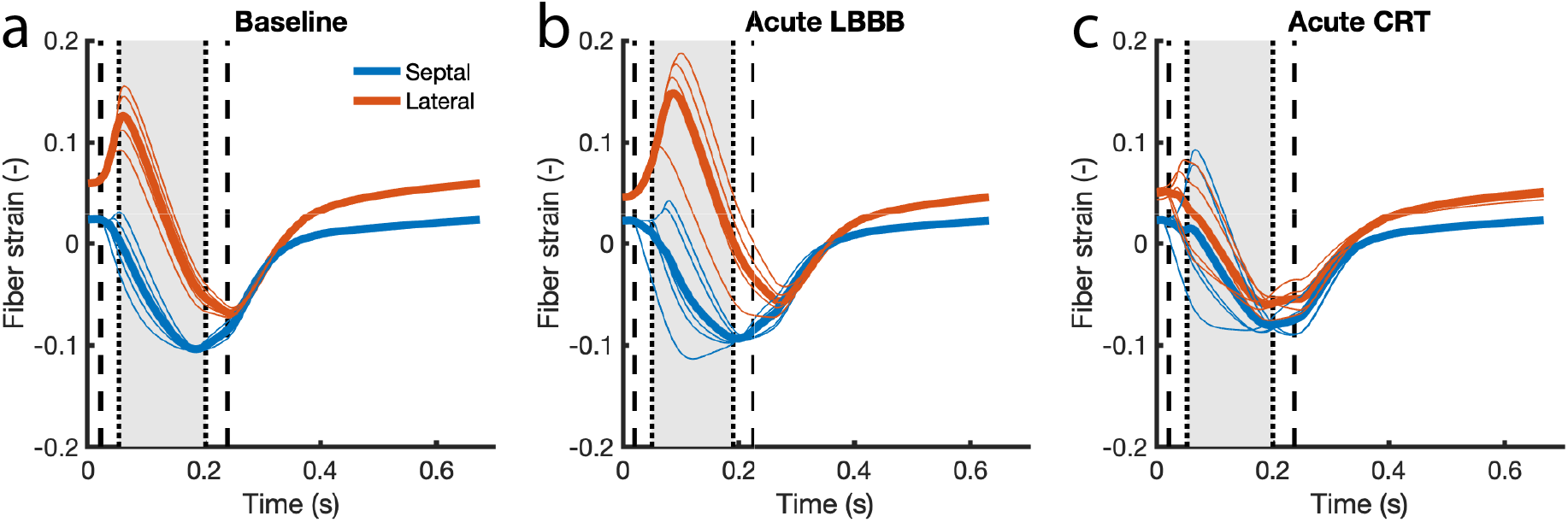
Mechanical dyssynchrony and maximum elastic fiber strain in late-activated segments, located in the lateral wall, increased from baseline to LBBB (b), and decreased following CRT (c). Average wall strain is shown by the thick lines, and valve opening and closure by black dashed (mitral valve) and dotted (aortic) lines.

The changes in maximum fiber strain gave rise to global and local changes in wall mass. During LBBB, LV mass increased (Fig. 4a), with growth mainly occurring in the late-activated lateral wall segments (Fig. 4b,c). Simulated CRT led to a reversal of these effects: peak strains in the lateral wall segments were reduced towards baseline due to earlier activation, causing these segment masses to regress. Although septal wall mass remained within the reported error bars during CRT, the model predicted a rapid increase early during CRT that was not observed in the experimental data. As a consequence of differences in activation timing and mechanically-driven growth, global LV function changed during LBBB and CRT. Acutely post-LBBB, stroke volume, maximum pressure and dp/dt_max_ decreased (Fig. 5a,b). During 8 weeks of LBBB, EDV and ESV progressively increased (Fig. 5c). Simulated CRT reversed these effects: stroke volume and dp/dt_max_ increased immediately after CRT, and EDV decreased towards baseline value over 8 weeks.

**Fig. 4.**
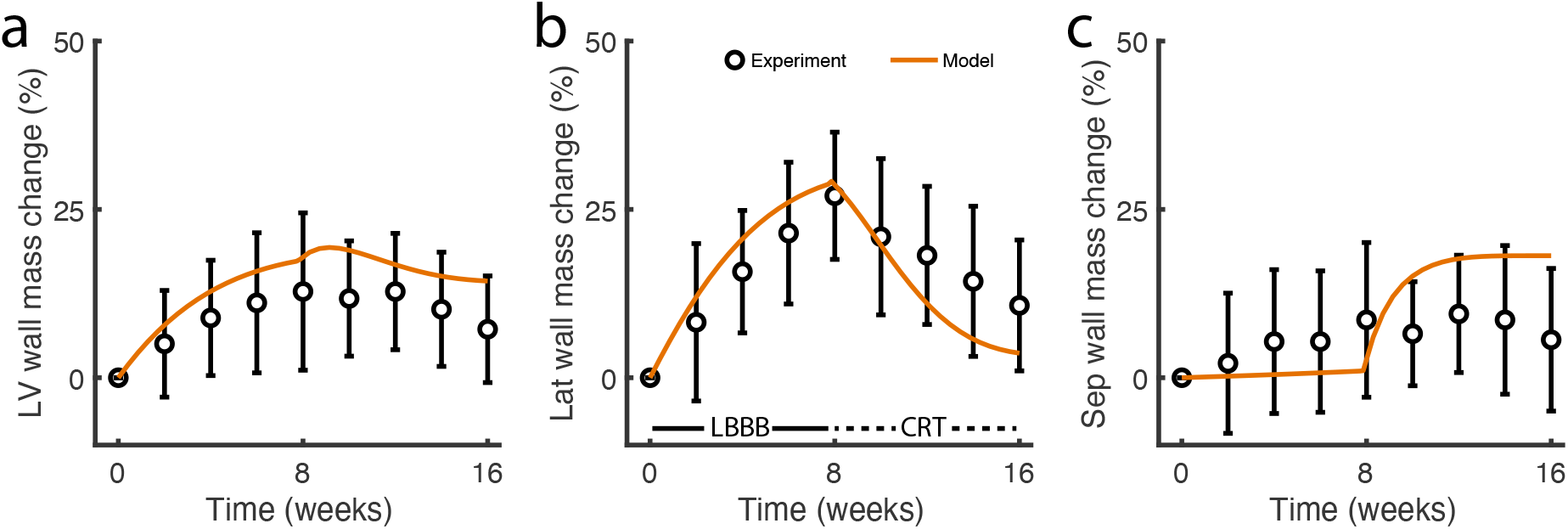
The remodeling predicted by the model fell within one standard deviation of the reported data for global (a), lateral (b), and septal (c) mass change during both LBBB (weeks 1-8) and CRT (weeks 8-16).

**Fig. 5.**
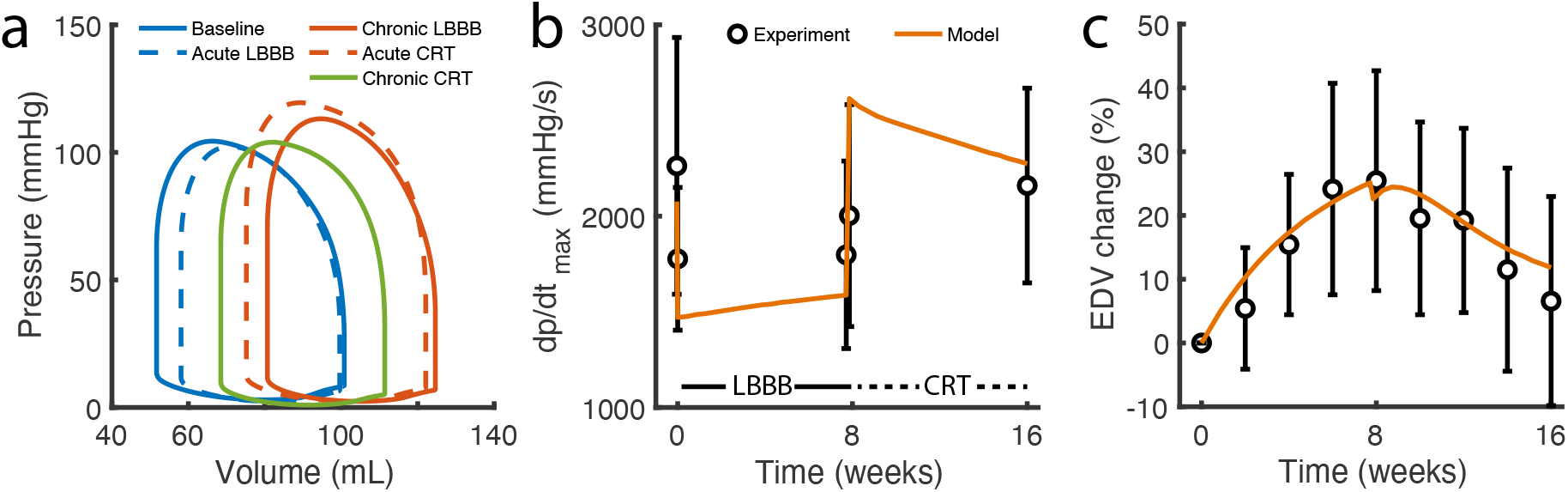
The model was calibrated to match reported hemodynamics. Following the introduction of LBBB, stroke volume (a) and dp/dtmax (b) decreased and EDV progressively increased (c). These effects were reversed during CRT.

### 3.3 LV pacing location influences post-CRT remodeling and is not correlated to acute outcome

With the calibrated model, we investigated the influence of LV lead location on chronic CRT outcome. After 8 weeks of LBBB, alternate CRT scenarios were simulated with the LV lead located in each of the 16 AHA segments, requiring less than 10 minutes total on a laptop computer. At the onset of CRT, all lead locations produced small decreases in EDV (3-5%, Fig. 6b); however, different lead locations led to dramatically different remodeling predictions. Some locations in the lateral wall produced up to 10% reductions in EDV and 30% reductions in lateral wall mass after 8 weeks of pacing, while other pacing locations produced little to no reversal of LBBB-induced dilation and hypertrophy.

**Fig. 6.**
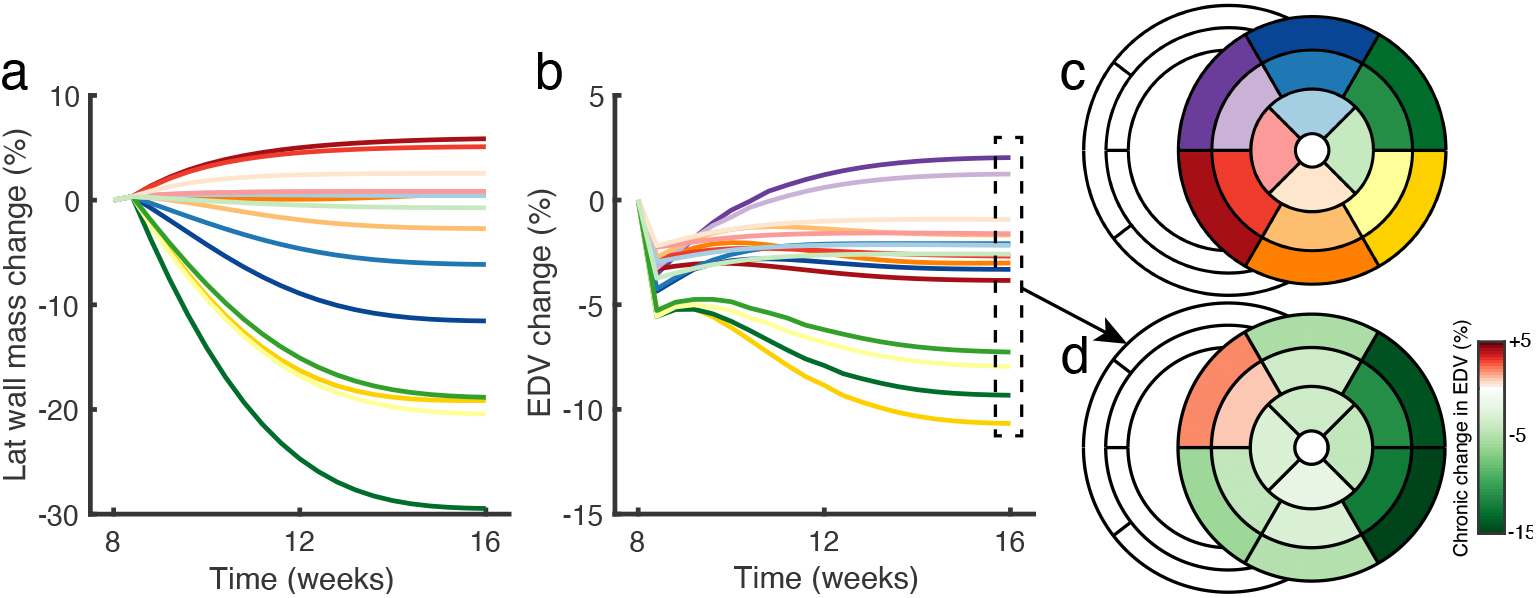
The calibrated model predicted that CRT remodeling outcome, i.e. reversal of lateral wall mass (a) and EDV (b), was dependent on pacing location. Pacing at the lateral basal wall segments resulted in the largest reduction of EDV (c,d) after 8 weeks of simulated CRT. The line colors in (a) and (b) match with the pacing location, as depicted in (c). Hemodynamics were assumed to remain constant during CRT.

For cases combining LBBB with different infarct locations, the lead location that resulted in the largest reduction in EDV, and the maximum amount of reverse remodeling that could be achieved depended on ischemia location (Fig. 7). The presence of (non-conductive and non-contractile) ischemia increased end-diastolic volume, and influenced the electrical conduction pattern, thus changing the magnitude and distribution of maximum strains, which drove remodeling in our model. Interestingly, and in contrast to the nonischemic LBBB prediction, the pacing site resulting in the best remodeling response for ischemic LBBB (except LAD midwall scar) did not correlate with the pacing site resulting to the best acute electrical response (shortening of QRS duration). Moreover, the predicted maximum amount of reverse remodeling achieved by CRT was different between ischemic cases, with the least amount of reverse remodeling predicted for LAD midwall ischemia (3%) and the most for LAD basal ischemia (14%).

**Fig. 7.**
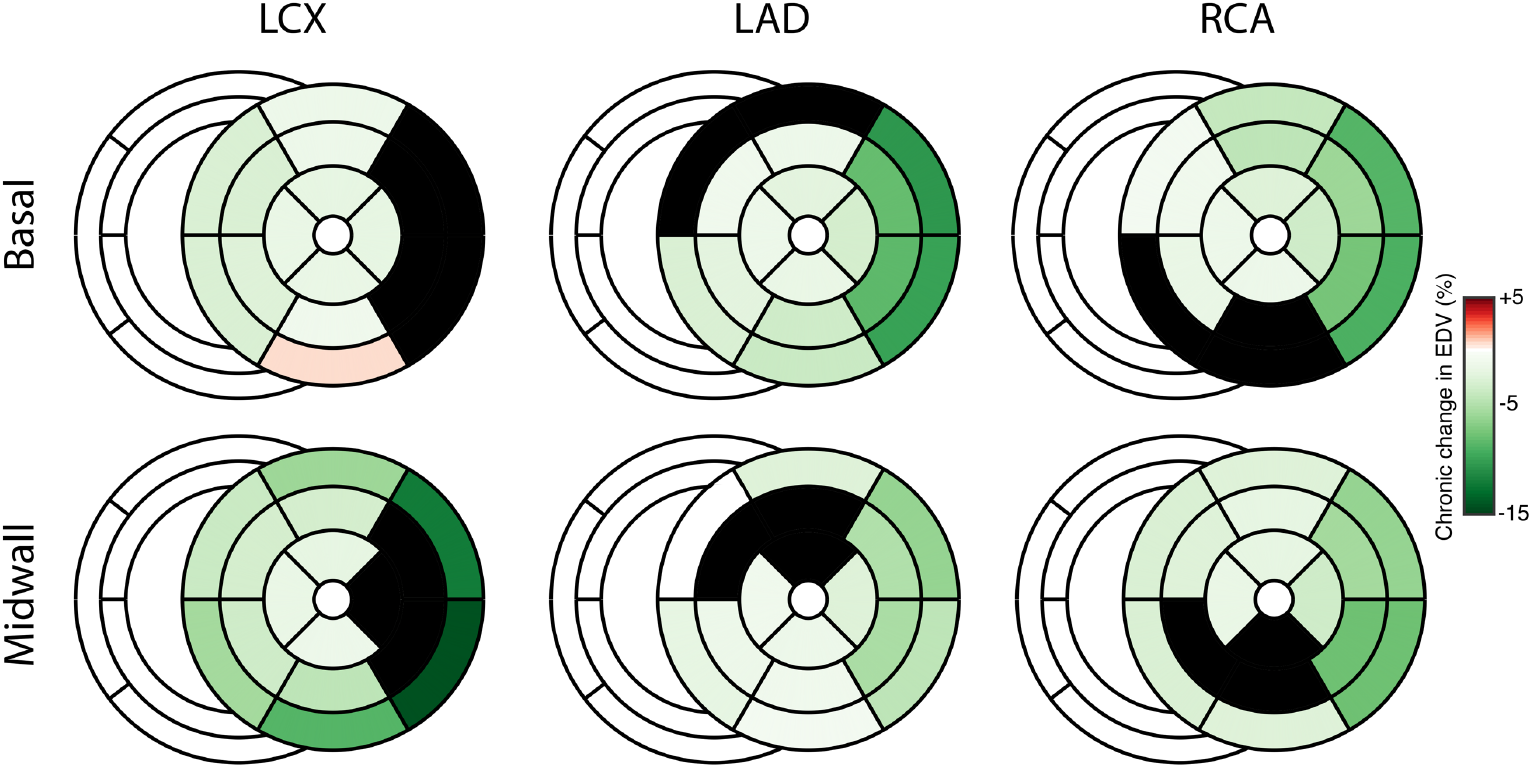
The model predicted that CRT remodeling outcome (8-week change in EDV) for ischemic LBBB cases is dependent on both pacing location and ischemia location (black segments).

## 4 Discussion

The goal of this study was to develop a rapid electro-mechanical model to predict reverse LV remodeling following CRT. We adapted our recently published rapid model of LV remodeling (Witzenburg and Holmes 2018) to simulate the mechanics and reverse remodelingfollowing ventricular dyssynchrony and CRT, and added a rapid electrical model to predict electrical activation timing. The model was calibrated to previously published experimental data (Vernooy et al. 2007) and applied to predict the influence of CRT lead placement on LV remodeling. Interestingly, the model predicted that for most ischemic LBBB cases that were simulated, short-term response to CRT (shortening of QRS duration) does not correlate with predicted long-term remodeling.

### 4.1 Rapid predictions of remodeling following CRT in a clinical setting

The long-term success of CRT is determined by reversing LV dilation (Bristow et al. 2004; Cleland et al. 2005; St. John Sutton et al. 2003). Currently, this desired response is only achieved in 50-65% of patients receiving CRT (Chung et al. 2008; Tracy et al. 2012). Therefore, we have developed a computational model to predict post-CRT remodeling. The model was tuned to accurately match experimental canine data of LV remodeling with heart failure LBBB and subsequently favorable remodeling with CRT (Figs. 4 5). It includes several features that were previously demonstrated to influence LV remodeling following CRT, such as activation timing at the lead position and scar location (Bilchick et al. 2014; Rademakers et al. 2010). As a result, it has the potential to predict patient-specific remodeling following CRT. Our initial predictions indicate that for many ischemic LBBB cases (Fig. 7), long-term remodeling outcome is not necessarily correlated to short-term change in QRS duration (Fig. 8). This implies that a pacing location that looks promising in the electrophysiology may not always be the optimal approach with respect to long-term outcomes. This finding is consistent with the clinical observation that the extent of CRT response tends to be lower in patients with ischemic LBBB St. John Sutton et al. (2003); Bilchick et al.(2014).

**Fig. 8.**
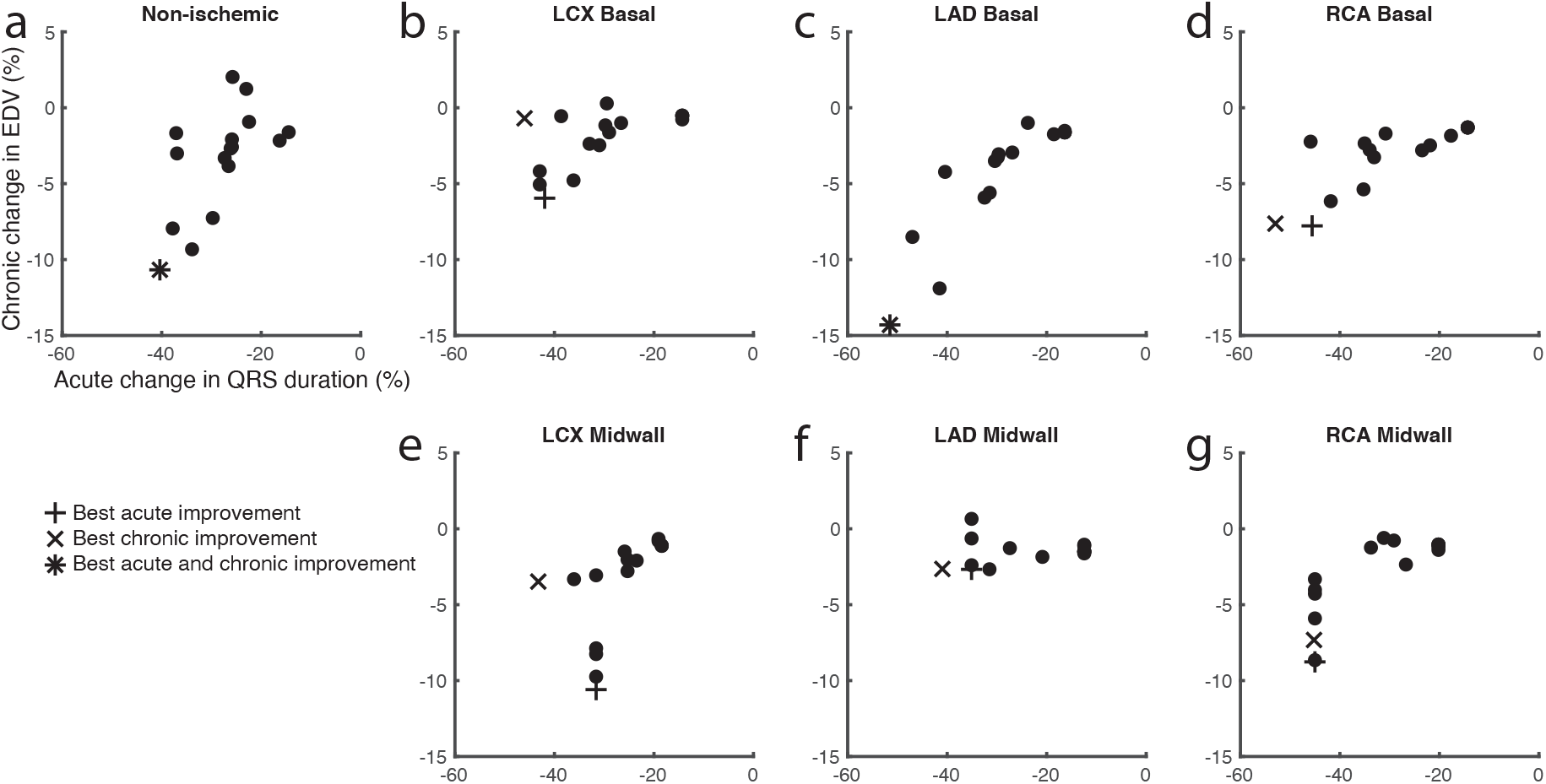
In non-ischemic LBBB (a), the CRT pacing location that was predicted to lead to the largest shortening of QRS duration (+) was also predicted to lead to the largest chronic reduction in EDV (×). However, including ischemia decoupled these two predictions for all ischemia locations except LAD basal (b-g).

A key feature of this model is that it is computationally efficient enough to allow the exploration of many different lead locations and pacing scenarios for each patient as part of the routine pre-procedure workflow, or even in near-real-time in the electrophysiology lab. This is due to the simplified ventricular geometry and fast rule-based model of electrical propagation. Here, a sweep of 8-week post-CRT remodeling predictions for lead locations at all 16 AHA segments took only 6 minutes on a laptop computer. The influence of additional CRT parameters on remodeling can thus be assessed in a reasonable time frame, such as different pacing modes and pacing timing of the LV lead. Note that the effect of atrioventricular delay can currently not be tested, since both our electrical and mechanical model lack atria. The computational efficiency of the model also allows for customizing this model to individual patients using our fitting method. This is important since medical data required to customize the model to individual patients, such as magnetic resonance imaging, echocardiography, or blood pressure measurements, are often collected as late as one day before the CRT procedure. We previously demonstrated that our model fitting method yields a unique solution without requiring user intervention (Witzenburg and Holmes 2018).

### 4.2 Modeling remodeling following LBBB and CRT

We previously modeled LV growth using a phenomenological growth relationship through which fiber and radial growth were independently driven by changes in maximum fiber and minimum radial strain, respectively (Witzenburg and Holmes 2018). This relationship was based on work from Kerckhoffs et al. (2012a), which was found to be most promising among multiple published growth models for predicting growth induced by both volume and pressure overload (Witzenburg and Holmes 2017). Applying this growth relationship here for dyssynchrony-induced growth did not achieve results that matched with experimental data. Following LBBB, minimum radial strain (inversely proportional to fiber strain, Fig. 3) did not increase in late-activated wall segments and the model therefore did not predict wall thickening, in disagreement with experimental observations (Vernooy et al. 2005; Prinzen et al. 1995; Vernooy et al. 2007). Therefore, we used a similar approach to Arumugam et al. (2019) and used maximum fiber strain to drive growth in both the circumferential and radial directions. Previous work by Arumugam et al. was based the formulation of the stimulus function and growth tensor on experimental findings that stretching cardiomyocytes longitudinally can induce addition of sarcomeres both in series and in parallel. This, in turn, results to growth in parallel and perpendicular to the stretching direction (Yang et al. 2016). The choice of isotropic growth was validated by matching both experimental increases in EDV (Fig. 5c), mainly determined by fiber growth, and wall mass (Fig. 4), determined by both fiber and radial growth. Additionally, the growth law was here augmented with an evolving homeostatic setpoint, which Yoshida et al. (2019) recently demonstrated to be key to predict reverse growth, as occurs following relief of pressure overload or during CRT.

### 4.3 Level of anatomical detail required to model LV strain during dyssynchrony

Modeling the mechanical behavior of ventricular wall segments is crucial to accurately remodeling using a strain-driven growth law. In our previous computational study of LV remodeling, left and right ventricular geometry were modeled as thin-walled spheres with the entire wall contracting simultaneously. In order to model ischemia, we used the approach of Sunagawa et al. (1983), in which the spherical LV compartment is functionally divided into multiple smaller spherical subcompartments. In this approach, subcompartments can have different wall stiffness and volume, which add up to total LV volume, but share the same pressure throughout the cardiac cycle. We found this approach to be adequate when predicting post-infarction remodeling using a non-contracting ischemic and a healthy subcompartment (Witzenburg and Holmes 2018, 2019). However, when we attempted to divid the LV into 16 spherical subcompartments according to the 16-segment AHA model, with onset of contraction of each subcompartment given by the electrical model, the model overestimated mechanical dyssynchrony during baseline and underestimated changes in strain between baseline and LBBB. Additionally, no reduction in pumping function following LBBB was achieved.

In the present study, we hypothesized that excess mechanical dyssynchrony at baseline using a Sunagawa-type approach was caused by the absence of spatial constraints between LV subcompartments. In the Sunagawa approach, LV subcompartments do not have a spatial location and therefore adjacent subcompartments can deform independently from one another. By contrast, in finite element models adjacent elements share common nodes such that all elements are connected and differences in deformation between adjacent elements are limited. We adopted the ventricular geometry of Lumens et al. (2009) as a middle ground that provides additional constraints on deformation of adjacent elements while preserving the speed of a compartmental approach. Lumens and coworkers model the ventricular geometry as three thick-walled spherical segments that encapsulate the two ventricular cavities, where each wall can be divided into multiple segments (Walmsley et al. 2015) that can have different wall stiffness due to local differences in electrical activation timing but all share the same curvature throughout the cardiac cycle, thus providing more geometrical stability. Walmsley et al. (2015) previously demonstrated that this model setup can accurately simulate post-LBBB mechanics, which is crucial to correctly predict strain-driven remodeling. In a subsequent study, and in contrast to our experience using the multicompartmental Sunagawa approach, Walmsley et al. (2016) showed that this model setup can also reproduce septal flash and septal rebound stretch, associated with reduction in LV pump function in patients with LBBB (Grines et al. 1989). In the work presented here, septal rebound stretch was not present (Fig. 3b), which may have caused the relatively poor match to Vernooy’s data on septal growth (Fig. 4b-c). However, Walmsley et al. (2016) found septal rebound stretch is dependent on factors including intraventricular differences in activation timing between septal and lateral wall. Because we calibrated our electrical model to our own data and not to Vernooy et al. (2007), intraventricular activation timing could have been different between the two studies.

### 4.4 Limitations

The current model has three main limitations. First, the phenomenological growth model was chosen and calibrated specifically for cardiac remodeling following LBBB and CRT in canines. This growth model may need to be reparameterized when applying it to predict human post-CRT remodeling. Moreover, the isotropic growth tensor may not be applicable to model growth that is caused by different pathologies, such as pressure and volume overload, as discussed above. Thus, any clinical application of this model should be preceded by retrospective calibration and prospective validation on an appropriate sample of patients representing the exact population for which it will be employed.

Second, the degree of mechanical dyssynchrony and maximum strain during baseline (Fig. 3) in our model is higher than reported in experimental studies (Prinzen et al. 1995; Vernooy et al. 2005; Aalen et al. 2019; Fixsen et al. 2019; Vernooy et al. 2007). This could be related to how we represented coupling between electrical and mechanical activation in the active contraction model. Kerckhoffs et al. (Kerckhoffs et al. 2003) similarly found in a modeling study that physiological depolarization resulted in less physiological strain patterns, suggesting that there is some local mechanism that makes contraction more synchronous than depolarization by variation of local electromechanical delay time. This hypothesis is supported by a subsequent modeling study by Constantino et al. (Constantino et al. 2012), who found that electromechanical delay increased in dyssynchronous heart failure compared to normal hearts. Nevertheless, in our model the difference in maximum strain between baseline and LBBB, and between early and late-activated segments during LBBB, were sufficient to accurately reproduce the local remodeling behavior following LBBB and CRT.

Third, our model does not predict hemodynamic regulation, which is known to play an important role in LV remodeling following pathologies such as pressure overload, volume overload, and myocardial infarction (Witzenburg and Holmes 2018, 2019; Rondanina and Bovendeerd 2020a,b). Instead, the model currently relies on being fitted retrospectively to experimental data, which limits its predictive capability. Since no experimental data was available to compare to our predictions of post-CRT remodeling for ischemic LBBB, we therefore decided to keep the hemodynamic parameters constant during those simulations. In the future, hemodynamic regulation mechanisms such as baroreflexes and renal function could be added to the model to regulate heart rate and hemodynamic parameters (capacitances, resistances, and SBV), as for example done by Beard et al. (2013) and Rondanina and Bovendeerd (2020a,b).

### 4.5 Conclusions

We developed and validated a fast electro-mechanical model to predict LV remodeling following CRT, then used the model to explore the impact of post-infarction ischemia on the ventricular remodeling produced by simulated CRT using different lead locations. For most ischemic LBBB cases we simulated, short-term response (shortening of QRS duration) did not correlate with long-term reverse remodeling. This finding is consistent with the clinical observation that the extent of CRT response tends to be lower in patients with ischemic LBBB. It also highlights the contributions a computational model could make to improving patient-specific customization of CRT in such patients. While this work represents just the first step towards predicting patient-specific CRT responses, we believe both the results and the time frame required to customize and run this model suggest promise for this approach in a clinical setting.

## Acknowledgements

This study was funded by the National Institutes of Health (U01 HL127654) and the Seraph Foundation.

