## Supplementary Data for "A Rapid Electromechanical Model To Predict Reverse Remodeling Following Cardiac Resynchronization Therapy"

**Noname manuscript No.**
(will be inserted by the editor)

### A Rapid Electromechanical Model To Predict Reverse Remodeling Following Cardiac Resynchronization Therapy

#### Supplementary Data

**Pim J.A. Oomen · Thien-Khoi N. Phung · Kenneth C. Bilchick · Jeffrey W. Holmes**

Received: date / Accepted: date

##### Supplement

Mechanical and growth model pseudocode

1. Provide initial estimate of compartmental volumes (blood vessels and ventricles) at cardiac cuycle time $t = 0$.
2. Estimate compartmental volumes at cardiac cycle time step $t$ via 4th order Runge-Kutta method
3. Calculate ventricular strain
   (a) Use converged solution of previous time step as initial estimate of junction circle radius and SW position
   (b) Calculate segmental strains and stress
   (c) Calculate wall tension
   (d) Check for balance of tension, if not satisfied update estimates of junction circle radius and SW position via Newton-Raphson method and repeat b an c.
4. Calculate compartment pressures
5. Repeat 2-4 for all cardiac cycle time steps
6. Check for steady-state solution, if the compartmental volumes at the beginning and end of the cardiac cycle were not within 0.1 mL of each other, update the compartmental volumes at the beginning with those at the end and repeat 2-5.
7. Determine segmental growth $F_g^{i+1}$ using elastic segmental strain, and update wall mass and unloaded midwall surface area. Repeat 1-7 for each growth step $i$.

Model parameters

v All model parameters are listed in Supplementary Table 1. For the electrical model, myocardial fiber conduction velocity $v_f$ was fitted to in-house 12-lead ECG data, as described in detail in Section 2.5. Parameters of the mechanical model (including constitutive model parameters) were fitted to data from Vernooy et al. (2005, 2007) using our previously published fitting method (Witzenburg and Holmes 2019b). Model parameters at pre and acute post-LBBB were fitted by minimizing the following objective function consisting of Z-scores of hemodynamic and kinematic measures (see Section 2.5):

$$E = \left(\frac{EDP^{pre}_{model} - EDP^{pre}_{exp}}{SDEDP^{pre}_{exp}}\right)^2 + \left(\frac{EDP^{post}_{model} - EDP^{post}_{exp}}{SDEDP^{post}_{exp}}\right)^2 + \left(\frac{ESP^{pre}_{model} - ESP^{pre}_{exp}}{SDESP^{pre}_{exp}}\right)^2 + \left(\frac{ESP^{post}_{model} - ESP^{post}_{exp}}{SDESP^{post}_{exp}}\right)^2 + \left(\frac{max\,(dp/dt)^{pre}_{model} - max\,(dp/dt)^{pre}_{exp}}{SDmax\,(dp/dt)^{pre}_{exp}}\right)^2 + \left(\frac{max\,(dp/dt)^{post}_{model} - max\,(dp/dt)^{post}_{exp}}{SDmax\,(dp/dt)^{post}_{exp}}\right)^2 + \left(\frac{EDV^{pre}_{model} - EDV^{pre}_{exp}}{SDEDV^{pre}_{exp}}\right)^2 + \left(\frac{EF^{pre}_{model} - EF^{pre}_{exp}}{SDEF^{pre}_{exp}}\right)^2 + \left(\frac{\varepsilon\, cor^{pre}_{model} - \varepsilon\, cor^{pre}_{exp}}{0.5}\right)^2 + \left(\frac{\varepsilon\, cor^{post}_{model} - \varepsilon\, cor^{post}_{exp}}{0.5}\right)^2$$

The growth parameters were fitted to match experimental changes over 16 weeks of EDV, global wall mass, and lateral and septal wall mass. While adjusting the growth parameters, experimental EDP and ESP were matched at chronic (8 weeks) LBBB, acute post-CRT, and chronic (8 weeks) CRT by adjusting SVR and SBV. We assumed SVR and SBV changed linearly in between the reported time points.

**Table 1** Model parameters

| Model | Parameter | Value | Unit | Source | Description |
|---|---|---|---|---|---|
| Electrical | $v_{\mathrm{f}}$ | $0.85^a$, $1.85^b$ | m/s | Fitted | Myocardial fiber conduction velocity |
| | $v_{\mathrm{purkinje}}$ | 600 | % | Constant | Relative purkinje fiber conduction velocity |
| | $v_{\mathrm{cf}}$ | 40 | % | Constant | Relative cross-fiber conduction velocity |
| Mechanical | SVR | $0.50^a$, $0.65^b$, $0.80.^c$ $0.75^d$, $0.65^e$ | mmHg·s$^-$1·mL | Fitted | Systemic vascular resistance |
| | SBV | $379^a$, $330^b$, $335^c$, $400^d$, $335^e$ | mL | Fitted | Stressed blood volume |
| | HR | $89^a$, $95^b$, $82^c$, $90^d$, $92^e$ | min$^{-1}$ | Vernooy et al. (2007) | Heart rate |
| | $V_{\mathrm{LFW}}$ | 7.33 | mL | Measured | Left free wall volume |
| | $V_{\mathrm{RFW}}$ | 5.54 | mL | Measured | Right free wall volume |
| | $V_{\mathrm{SW}}$ | 4.67 | mL | Measured | Septal wall volume |
| | $A_{\mathrm{m,LFW}}$ | 81.8 | cm$^2$ | Fitted | Reference left free wall midwall surface area |
| | $A_{\mathrm{m,RFW}}$ | 105.5 | cm$^2$ | Fitted | Reference right free wall midwall surface area |
| | $A_{\mathrm{m,SW}}$ | 52.5 | cm$^2$ | Fitted | Reference septal wall midwall surface area |
| Constitutive (passive) | $k_1$ | 59.7 | kPa | Fitted | Passive behavior linear parameter |
| | $k_3$ | 0.57 | kPa | Fitted | Exponential scaling parameter |
| | $k_4$ | 25 | - | Constant | Exponential parameter |
| Constitutive (active) | $\sigma_a$ | 150.0 | kPa | Fitted | Maximum active stress |
| | $L_{\mathrm{s,ref}}$ | 2.0 | μm | Lumens et al. (2009) | Reference sarcomere length |
| | $L_{\mathrm{sc,0}}$ | 1.51 | μm | Lumens et al. (2009) | Contractile element length at zero stress |
| | $L_{\mathrm{se,iso}}$ | 0.04 | μm | Lumens et al. (2009) | Series elastic element length |
| | $v_{\mathrm{max}}$ | $5.0 \cdot 10^3$ | μm·ms$^{-1}$ | Kerckhoffs et al. (2012) | Maximum sarcomere shortening velocity |
| | $T_{\mathrm{act}}$ | 168 | ms | Constant | Base duration of (length-dependent) contraction |
| Growth | $F_{\mathrm{g,max,pos}}$ | 0.1 | - | Fitted | Maximum forward growth rate |
| | $F_{\mathrm{g,max,neg}}$ | 0.03 | - | Fitted | Maximum reverse growth rate |
| | $s_{\mathrm{50,pos}}$ | 0.075 | - | Fitted | Stimulus value at 50% of $F_{\mathrm{g,max,pos}}$ |
| | $s_{\mathrm{50,neg}}$ | 0.110 | - | Fitted | Stimulus value at 50% of $F_{\mathrm{g,max,neg}}$ |
| | $n_{\mathrm{pos}}$ | 3 | - | Fitted | Exponential parameter of forward growth |
| | $n_{\mathrm{neg}}$ | 9 | - | Fitted | Exponential parameter of reverse growth |

[a]Baseline, [b]Acute post-LBBB, [c]Chronic post-LBBB, [d]Acute post-CRT, [e]Chronic post-CRT

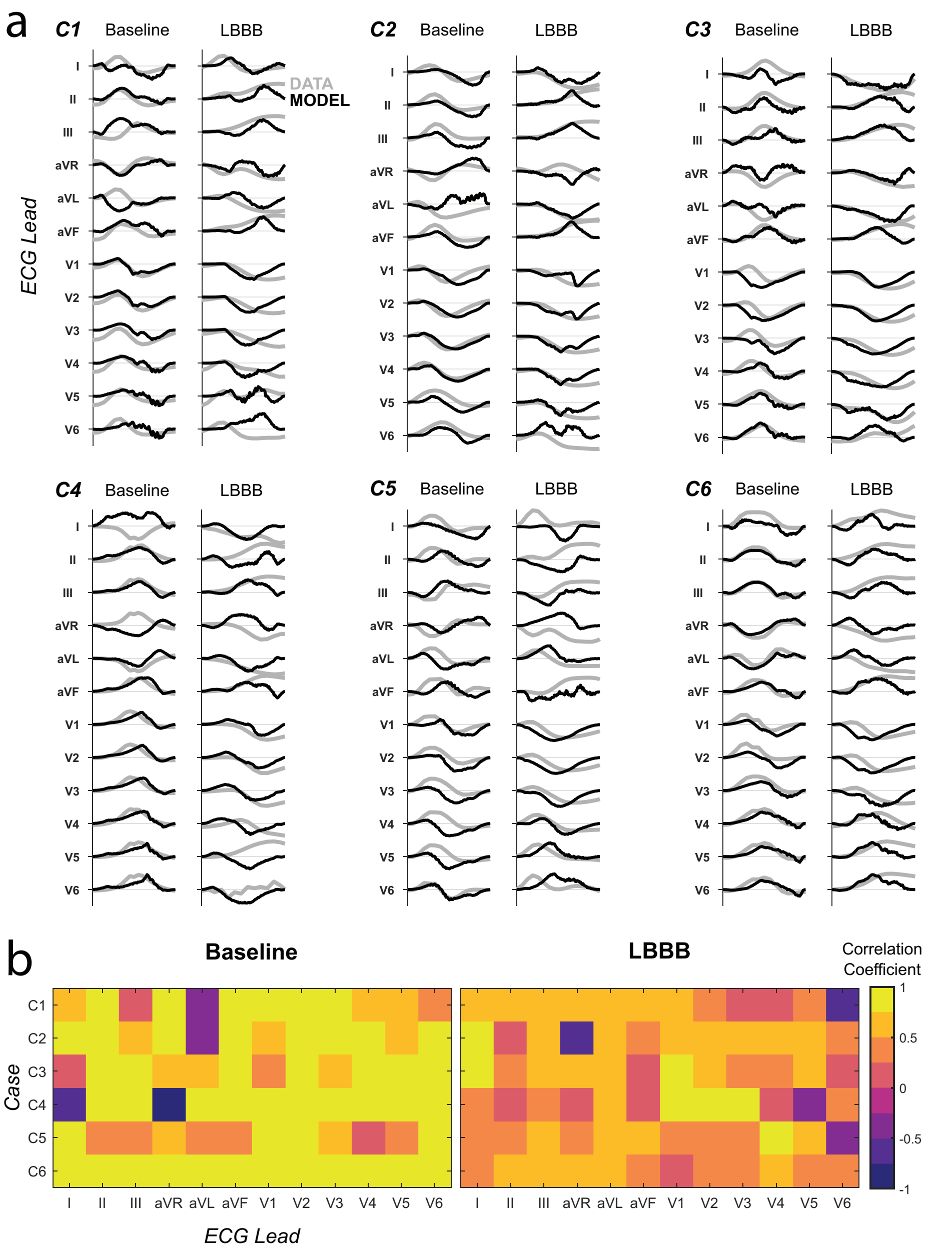


**Supplementary Figure 1:** Electrical model fits for all six canine data sets (C1-2 ischemic, C3-6 non-ischemic). (a) The best fit model pseudo-ECG is shown on top of the 12-lead ECG data. The ranges for the signals are normalized to compare signal shapes. (b) The correlation coefficients between model and data signals for each lead are shown in the heat map. C6 is the case used for all growth simulations.
